# BertNDA: A Model Based on Graph-Bert and Multi-scale Information Fusion for ncRNA-disease Association Prediction

**DOI:** 10.1101/2023.05.18.541387

**Authors:** Zhiwei Ning, Jinyang Wu, Yidong Ding, Ying Wang, Qinke Peng, Laiyi Fu

## Abstract

Non-coding RNAs (ncRNAs) are a class of RNA molecules that lack the ability to encode proteins in human cells, yet play crucial roles in various biological process. Understanding these relationships and how different ncRNAs interact with each other to affect diseases can vastly contribute to their diagnosis, prevention, and treatment. However, predicting tertiary interactions between ncRNA-disease associations by utilizing structural information across multiple scales remains a challenging task. It should be noted that research on predicting tertiary interaction between trinary ncRNA-disease associations is scarce, highlighting the need for further studies in this area. In this work, we propose a predictive framework, called BertNDA, which aims to predict association between miRNA, lncRNA and disease. The framework employs Laplace transform of graph structure and WL (Weisfeiler-Lehman) absolute role coding to extract global information. Local information is identified by the connectionless subgraph which aggregates neighbor feature. Moreover, an EMLP (Element-wise MLP) structure is designed to fuse the multi-scale feature representation of nodes. Furtherly, feature representation is encoded by using a Transformer-encoder structure, the prediction-layer outputs the final correlation between miRNA, lncRNA and diseases. The 5-fold cross-validation result furtherly demonstrates that BertNDA outperforms the state-of-the-art method in predicting assignment. Furthermore, an online prediction platform that embeds our prediction model is designed for users to experience. Overall, our model provides an efficient, accurate, and comprehensive tool for predicting ncRNA-disease associations. The code of our method is available in: https://github.com/zhiweining/BertNDA-main.

## I. Introduction

NON-CODING RNAs (ncRNAs) are RNA molecules that do not code for proteins but have essential roles in various biological processes in the human body[1]. Long-noncoding RNAs (lncRNAs), which are over 200 nucleotides long, form the largest class of ncRNAs and play critical roles in transcription, translation, splicing, epigenetic regulation[2-4]. Certain lncRNAs such as HOTAIR and UCA1 can serve as potential biomarkers for hepatocellular carcinoma recurrence and bladder cancer diagnosis[5, 6]. MiRNAs are another major class of ncRNAs that typically repress gene post-transcriptionally by binding to target mRNA’s 3’-untranslated regions (UTRs)[7]. Mounting evidence suggests that miRNAs significantly affect cell development, proliferation, and differentiation[8-10]. Meanwhile, some studies show that lncRNAs and miRNAs regulate each other[11], for example, lncRNAs can act as miRNA sponges, reducing their regulatory effect on mRNAs. Likewise, miRNAs bind lncRNAs and control stability of LncRNA[12]. Taken together, these findings underscore the importance of ncRNAs in the modulation of gene expression and their potential implications on human disease development. Therefore, it is necessary to further investigate the relationships between different ncRNAs and diseases, as well as the intrinsic affect among different types of ncRNAs (tertiary interactions) for better understanding the underlying mechanisms and develop effective therapeutic interventions.

In the past, it has been proven that studying associations in biomedical networks can uncover potential and biologically meaningful relationships including lncRNA-disease associations (LDAs)[13] and miRNA-disease associations (MDAs)[14, 15]. Although the traditional methods based on biological experiments achieve high accuracy, they often suffer from time-consuming and inefficient process. To address the issue of selecting prediction objects blindly, various machine learning algorithms have been developed to provide effective strategies for potential correlation prediction.

However, predicting tertiary interactions between ncRNA-disease associations remains a challenging task due to the complex structural information involved at multiple scales. The current research field in the bilateral relationship between RNA and disease is relatively mature, and we can extend existing methods to trilateral prediction relationships. One such traditional machine learning method, PADLMHOOI[16], implements high-order orthogonal iterations to predict potential associations and evaluates predictive performance through leave-one-out cross-validation. DFELMDA[17] employs a novel computational approach of random forest ensemble learning to predict miRNA-disease associations, integrating autoencoders for low-dimensional feature representation. Chen et al[18] postulate that functionally similar lncRNAs may be associated with similar diseases, and introduce the Laplacian regularized least squares method (LRLSLDA) within a semi-supervised learning framework to match lncRNAs with diseases. Although these traditional machine learning methods have demonstrated encouraging results, their prediction efficiency may decrease with increasing data volume, resulting in poor prediction performance for deeper correlations. Due to the relatively simple structure of these algorithms, the acquisition of multi-hop connection relationships in biological networks is not profound enough.

The difficulty of predicting tertiary interactions arises from the intricacies of understanding how different ncRNAs interact with each other and with diseases. Compared with the general tensor-based method, the graph structure has a more concise topological representation, and the edge information can obtain the potential connection relationship more accurately. Therefore, the graph structure has also received certain attention utilizing in the field of bioinformatics. In particular, GCNLDA[19] is a graph-based approach that constructs a heterogeneous graph comprising lncRNAs, diseases, and miRNAs. Then, the graph structure is utilized to learn local representation of lncRNA-disease pairs through graph convolutional networks and convolutional neural networks. GATGCN[20] is another method that employs both graph attention networks and graph convolutional networks to detect potential correlations between circRNAs and diseases. HGCNMDA[21] incorporates a gene layer in constructing a heterogeneous network. The model refines the feature of nodes into initial features and induction features and then learns miRNA and disease embeddings via a multi-graph convolutional network model. There are also some methods utilized attention mechanism to obtain a larger scale of feature representation in the process of molecular information. LDAformer[22] leverages topological feature extraction and Transformer encoder. Specifically, a pivotal process is designed for extracting potential multi-hop path feature from adjacent matrices during the topological information extraction stage. More recently, Zhan et al propose an approach called Graph-BERT[23], which is employed for node’s classification task. The method is trained through linkless subgraphs in the local context. Nevertheless, one limitation of the approach is the insufficiency to access global feature.

Despite the current models have achieved satisfied results, there are still many deficiencies, including only predicting the association between ncRNAs and diseases but ignoring the interaction between different ncRNAs. Meanwhile, underutilization of features within the associated graph structure of entities causes information ambiguity. To address these inadequacies, we propose a method named BertNDA based on Graph-BERT and multi-scale information fusion. Firstly, by collecting various biological datasets, the bioinformatics heterogeneous network including miRNAs, lncRNAs, and diseases is constructed. By calculating similarity between different molecules based on correlation information, it is able to extract information kernels for both functional and Gaussian interaction profiles. An EMLP module is used to extract global information from nodes using the Laplace transform of graph structure and WL absolute role embedding. Profiting from the design, the calculate cost of the backbone network can be rapidly decreased and the global feature can be aggregated in a more efficient way. Additionally, we calculate local information through a connectionless subgraph of nodes to avoid alleviating extract feature. Finally, multi-scale information fusion yield node characteristic representation with global self-attention mechanism employed as backbone of our model. Experimental results demonstrate that this method can furtherly reveal potential correlations at a global receptive field. A prediction-layer carry out the output of the correlation value, which effectively promotes people’s understanding of the basic level of complex diseases and assists the research of basic pathological molecules. The main contributions of the method are summarized as follows:

- Efficient prediction for trilateral interactions between ncRNA-disease associations: Compared with the previous bilateral prediction studies with ncRNAs and diseases, we supply the prediction of the relationship between ncRNAs, forming a more comprehensive trilateral prediction relationship, which is helpful for the study of molecular pathology.
- Global information representation utilizing full graph structure: In this study, we employ the heterogeneous graph structure to represent global features extracted from the Laplace matrix and WL absolute position coding.
- Local information aggregation from unconnected subgraphs: To obtain local information of nodes, we sort the neighbor nodes by intimacy and use the unconnected subgraphs to aggregate information about each node.
- Representation encoding using self-attention backbone network: We use the encoder part of the Transformer network as the backbone network of our method. By extracting the association relationship of nodes in a more macroscopic scale, we improve the effectiveness of the overall prediction.

## II. Materials

### A. Dataset

At present, a large number of biomedical experiments have confirmed that there is a certain causal relationship between many non-coding RNAs and diseases, and various biomedical websites also collect relevant experimental results, such as miRbase[24], HMDD[25], miRDisease[26], MNDR[27], LncRNADisease[28]. Making full use of these studies will be of great help to our research, and the databases acquired in this experiment are mainly from the following three aspects:

#### 1) miRNA-Disease associations

The miRNA-Disease associations we need are obtained from the two databases: (1) HMDD[25]: HMDD database is a manually collected and sorted miRNA-disease associations database. A total of 1206 miRNAs, 893 diseases, and 35547 pairs of associations are included. (2) miRCancer[29]: miRCancer currently documents 878 relationships between 236 microRNAs and 79 human cancers through the integration of >26000 published articles.

#### 2) lncRNA-Disease associations

The main sources of lncRNA-disease associations data we acquire are: (1) LncRNADisease[28]: the database integrates near 3000 lncRNA-disease pairs, including 914 lncRNAs and 329 diseases from about 2000 publications. (2) Lnc2Cancer3.0[30]: Lnc2Cancer3.0 database increases cancer-associated lncRNA entries over the previous versions. The release includes 9254 lncRNA-cancer associations, with 2659 lncRNAs and 216 cancer subtypes.

#### 3) miRNA-lncRNA associations

Currently, there is relatively little research on the interaction between miRNAs and lncRNAs, so we mainly rely on StarBase[31]: StarBase is designed for decoding the Interaction Networks of lncRNAs, miRNAs, competing endogenous RNAs(ceRNAs), RNA-binding proteins (RBPs) and mRNAs. In this section, we download two versions (2015 Version and 2017 Version) of lncRNA-miRNA associations datasets from starBasev2.0 database, which provides the comprehensive experimentally confirmed lncRNA-miRNA interactions based on large-scale CLIP-Seq data.

Due to the certain difference in the naming rules of diseases in various datasets, before data integrating, we preprocess the naming of various diseases to eliminate duplicate entities and make the adopted datasets more standardized. Then, by collecting the three categories of datasets, we obtain two datasets, named Dataset1 and Dataset2:

- Dataset1 is based on the three association relationships to find the intersection operation, that is: if a disease exists in both miRNA-disease and lncRNA-disease database, it will be included in dataset 1. The advantage of this method is that it can improve the density of the overall dataset, which makes great help for later prediction task. Finally, in dataset1, there are 7322 pairs of associations, including 4802 miRNA-disease associations, 2278 miRNA-lncRNA associations, 242 lncRNA-disease associations, and a total of 464 miRNAs, 56 lncRNAs, 121 diseases.
- Dataset2 is obtained by the union of the above three kinds of datasets, that is: if a disease is collected in either lncRNA-disease or miRNA-disease database, it will be included. The miRNAs and lncRNAs are approached in the same way. The amount of data can be significantly expanded by this manner. Finally, in Dataset 2, there are 43869 pairs of associations in the data obtained. The details are shown as Table 1 and experiments are mainly based on these two datasets for related design.

### B. Method

#### 1) BertNDA

To predict the potential associations in the tertiary entities, a method named BertNDA is proposed, which is based on Graph-BERT[23] model and multi-scale information fusion representation. Graph-BERT is a model that transfers BERT model to node classification tasks in the area of graph, and the information extraction of multi-scale features carried out on our method is applied to link prediction. At the same time, compared with other ncRNA and disease prediction models, our model performs excellent results in a variety of indicators.

The overall flow diagram of BertNDA is illustrated in Fig.1, which consists of four parts. Each part corresponds to the important component of the supervised prediction task. In the overall design process, we use the data structure of the graph to obtain a more complete molecular representation that is convenient for information extraction; At the same time, as an innovation of this study, the feature extraction part uses a multi-scale information representation combining global and local information, and also visualizes the extracted features in experiment section. With its superior structural performance, the global self-attention mechanism can be better at capturing the internal correlation of data, which is employed as the backbone network in our prediction task. Finally, the excellent prediction effect can be achieved based on the powerful representation ability of the model.

**Fig.1.**
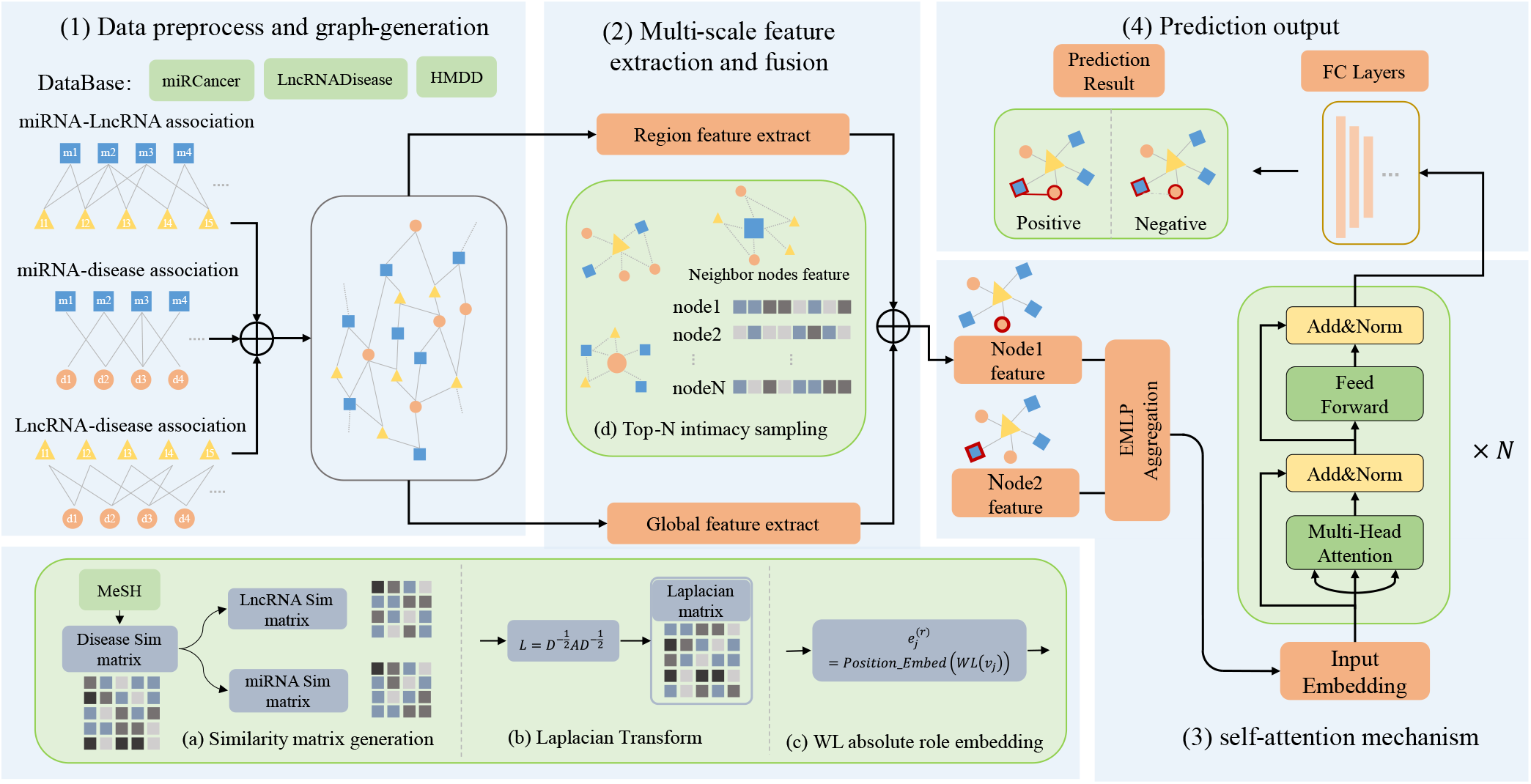
The flowchart of BertNDA. (1) Data preprocess and graph-generation part: the data sets of major official websites are collected to obtain the connection relationship between the two pairs and further transform it into a heterogeneous undirected tertiary graph; (2) Muti-scale feature extraction part: the global feature and local feature of the node are acquired, in which the global feature includes the similarity matrix, the Laplace matrix of the graph, the WL absolute position embedding, and the local feature is combined by the neighbor node feature in the subgraph; (3) Self-attention encoder part: after aggregating the feature by an EMLP module, the encoder structure of the Transformer is adopted as the main coding network; (4) Prediction output part: the encode result is passed through the output of the prediction layer to calculate the final result.

#### 2) Generation of Biomolecular Graph Similarity Matrix

Based on the relevant data collected and sorted, a multi-source biomolecular network 𝒢 = (*V*, ℰ) is constructed, where *V* represents the set of nodes, and ℰ represents the set of edges in the network. Moreover, the network corresponds to the association matrix is *A* ∈ *R*^*N*×*N*^(*N* = 641) in dataset1. If there is an experimentally verified relationship between a disease node *d*_*i*_ and a miRNA node *m*_*i*_, element *A*(*d*_*i*_, *m*_*i*_) is equal to 1 in the association matrix, otherwise, it is equal to 0.

##### 1) Disease semantic similarity

Since the naming of diseases is generally related to the symptoms they produce, the associations can be constructed according to the naming of diseases. Specially, MESH[32] is a unified database that uses a tree structure to characterize the correlation of diseases according to the naming, based on which we can calculate the disease semantic similarity. Supposing for a disease *d*_1_, *D*_1_ represents the set of *d*_1_ and all its ancestor diseases. Then, the semantic contribution value of any node *d*_*i*_ in the *D*_1_ to the *d*_1_ is defined as:

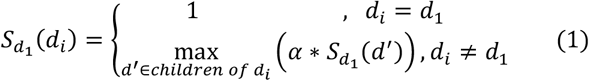

Where the *α* is an adjustable parameter, which we take the value of 0.5 according to previous work[17]. For any two diseases *d*_1_ and *d*_2_, their semantic similarity is defined as:

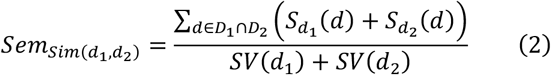

Where 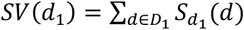 represents the sum of all the diseases’ semantic contribution in *D*_1_, so as *SV*(*d*_2_). It can be seen that the repetition in ancestral diseases between two diseases makes an influence in their semantic similarity. As a reliable way to judge the association of diseases, the semantic similarity significantly improves the efficiency of feature extraction.

##### 2) Disease Gaussian interaction profile kernel similarity

The external information on the relationship between diseases and ncRNAs can also be exploited. From these known associations, we can obtain the Gaussian interaction profile kernel similarity of diseases. For any two diseases, *d*_*i*_ and *d*_*j*_, the GIP similarity between them is calculated as follows:

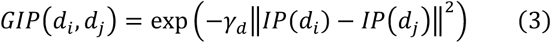

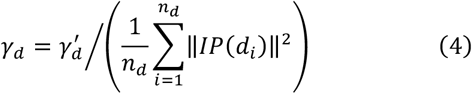

Where 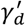 is the normalization parameter, and we set it to 1 according to previous work[18]. *n*_*d*_ indicates the total number of diseases. *IP*(*d*_*i*_) represents the vector corresponding to the *d*_*i*_ in the association matrix. The GIP similarity of diseases descripts that when the association vector *IP*(*d*_*i*_) of a disease *d*_*i*_ has a closer Euclidean distance to the association vector *IP*(*d*_*j*_) of another disease *d*_*j*_, the similarity score of the two diseases is also higher.

Considering that the semantic similarity between some diseases may be equal to 0, we combine the semantic similarity with GIP similarity to obtain the final similarity metric between diseases, which is defined as follows:

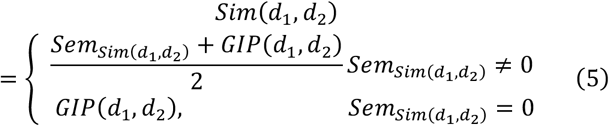

##### 3) Functional similarity of miRNA/lncRNA

Based on the hypothesis that ncRNAs with similar functions are more likely to cause the same disease[33]. We use disease similarity and the association matrix to obtain the functional similarity between ncRNAs, which is calculated as follows:

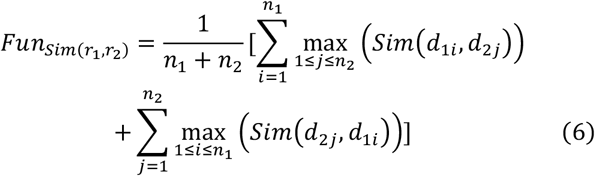

Where *d*_1*i*_(1 ≤ *i* ≤ *n*_1_) and *d*_2*j*_(1 ≤ *j* ≤ *n*_2_) indicate diseases associated with *r*_1_ and *r*_2_, respectively. And *Sim*(·) function indicates the similarity of diseases.

In summary, we acquire the semantic similarity and Gaussian interaction profile kernel similarity matrix between diseases, the functional similarity matrix between ncRNAs. The all similarity matrices between the constructed pairs are added to the original association matrix A. Then, the final similarity matrix *M*_*S*_ ∈ *R*^*N*×*N*^ is acquired.

#### 3) Global Feature Extraction and Fusion Based on EMLP

Global feature is a significant composition of the deep learning, which can quickly obtain key information from the entire data structure. In the field of graphs, it is challenging to design a unique global node representation because there are symmetries which prevent canonical node positional information[34]. This also makes some current graph networks based on attention mechanism, such as GAT, usually obtains the domain information of nodes by their neighbors[35]. In this work, we calculate the global information of the node in the following two ways:

##### 1) Laplace transform of graph structure

We construct a multi-source bioinformatics network containing three types of molecules, which is an undirected graph structure. In general graph structures, positional encoding is commonly used for feature representation of nodes. It is critical to ensure that each node has a distinct representation. Specially, Dwivedi[36] used the structure of the connectionless graph to acquire the Laplace matrix, and represented the Laplace feature vector of each node as the position. By this approach, nearby nodes have similar positional feature and farther nodes have dissimilar positional feature.

We calculate the Laplace matrix on the structure of the entire graph after data preprocessing. Eigenvectors are defined via the factorization of the graph Laplacian matrix, and its general definition and normalization are defined as follows:

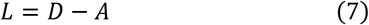

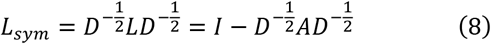

Where *D* represents the degree matrix 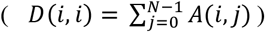, *A* represents the original association matrix. Each row 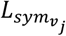 in 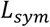 represents the Laplace feature vector of the node *v*_*j*_.

##### 2) Weisfeiler-Lehman Absolute Role Embedding

Besides, we also use WL (Weisfeiler-Lehman) absolute role representation[37] for further extraction of the global feature. The algorithm assumes that nodes with the semblable role in the graph have more similar code feature representation. And for two isomorphic graphs, the set of representation of the nodes is consistent. Therefore, the feature representation can be well embedded based on the relationship between nodes and graph.

The absolute role embedding of nodes is calculated by using the WL algorithm. Specifically, for each node *v*_*i*_, the method firstly acquires the definition set of the nodes connected to it (the initial definition of each node is set to 1). Secondly, the hash monotypic function is used to aggregate the features of the surrounding points set to the current node *hash* 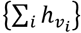. Then, the process is duplicated until the definition of each node is convergent. Finally, the WL eigenvalue *WL*(*v*_*i*_) about the node is obtained, then the eigenvalue is used to calculate the node feature embedding 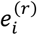, which is defined as follows:

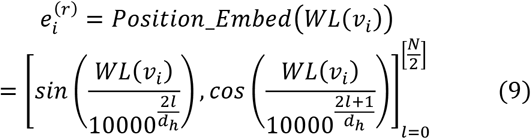

Where 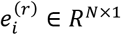, and *l* represents an index value, which is traversed from 0 to 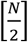. Similarity to the previous work[38], each element in the vector is computed by trigonometric functions sin(·) and cos(·).

##### 3) EMLP structure

Currently, there are two main ways to carry out feature aggregation in previous works, (i) one is to add multiple feature vectors after assigning different learnable weights; (ii) the other is to obtain feature vector with longer dimensions through direct splicing of vectors, and then feature dimensionality reduction is carried out through MLP. However, the EMLP(Element-weight MLP)[39] method is adopted to information aggregation in our work. The study shows that the feature information obtained by this way can remain the key element weights in the vector well compared with the first method and also has better learning effect and robustness than the second method. The flowcharts of these three methods are illustrated in Fig.2.

**Fig.2.**
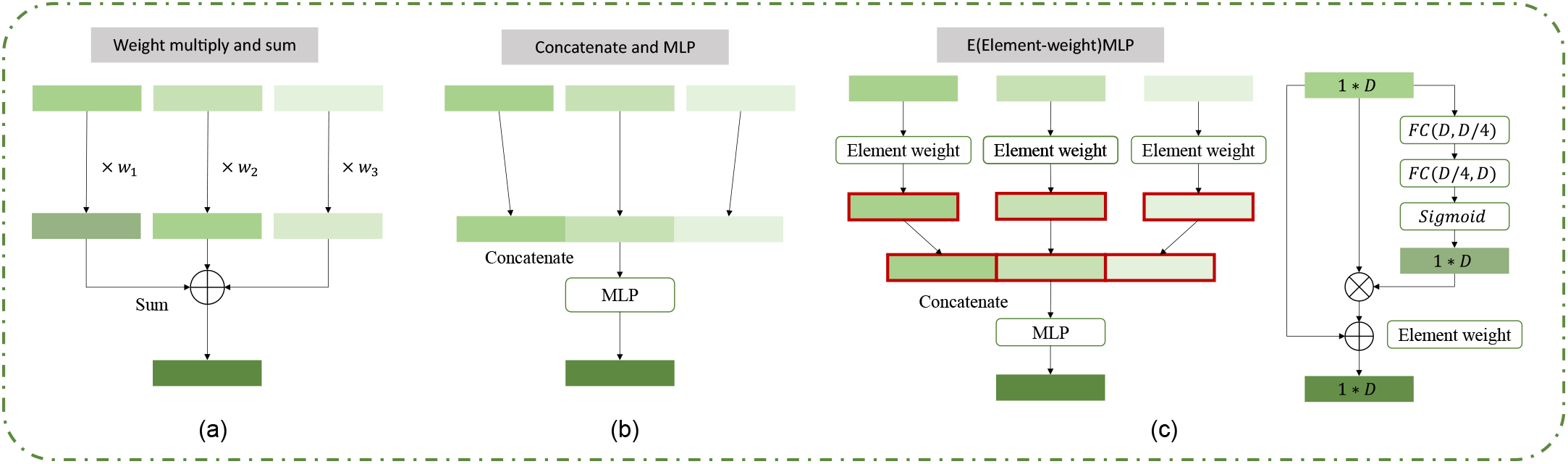
The three structures of feature aggregation. (a) Multiplied by the learnable weights and sum up; (b) Concatenate the features and employ normal MLP to reduce dimension; (c) The element-weight adopted in our method.

Concretely, our method learns the weight in element level for each feature vector *f*_*i*_, defined as follows:

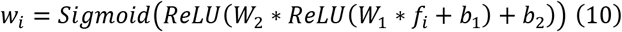

Then, the learned weight vector is multiplied with the origin feature vector, which can effectively remove the interference of other redundant information. After that, the newly obtained feature vector is added to the initial vector, defined as follows:

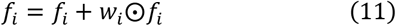

Where the operation ⨀ defines the element-wise product. Furtherly, the dimensionality reduction of information is carried out through a multilayer perceptron. Compared with directly combining the feature vectors into a matrix as the input into the backbone of subsequent predictions, the method of input after dimensionality reduction can decrease the calculation capacity and improve the efficiency of the overall model operation.

#### 4) Local Feature Extracted from the Unconnected Subgraph

In this section, we present the extraction process of local feature. Regional feature is considered as an important mean of extracting detailed information, which has more specific representation and can make full use of surrounding nodes’ information. In our study, we aggregate neighbor nodes by using a connectionless subgraph to gain local features. Firstly, we calculate the intimacy matrix between nodes[40], which is defined as:

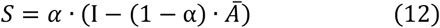

Where *Ā = AD*^−1^ represents the connection matrix after row normalization, *α* ∈ [0,1] and we set it to 0.15. For a node *v*_*i*_, we define its context nodes set as:

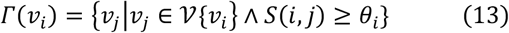

Where *θ*_*i*_ defines the intimacy threshold. Then, the top *k* nodes with the highest intimacy are selected to form the linkless subgraph *ℬ* together with *v*_*i*_, where ℬ ⊆ 𝒢. The corresponding feature vectors of *k* nodes are spliced together to generate the local feature matrix *M*_*L*_ ∈ *R*^*k*×*N*^ of the node *v*_*i*_.

#### 5) Backbone Network Based on Transformer Encode

The representation of node pairs are fused and utilized as the input to the Transformer encoder component to obtain the entity-pair associations encoding by self-attention mechanism. Concretely, the global feature vector of the combined nodes and their respective local feature matrix are acquired, which is deemed as the input to our backbone network. The output feature encoding is calculated using multi-heads attention mechanism and multi-layers self-attention encoding approach. The component reduces the feature dimension to a low-dimension, and the self-attention is calculated as follows:

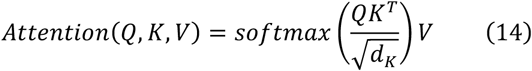

In each attention head, three matrix multiplications are used to convert X into Q, K, and V. The calculation is as follows:

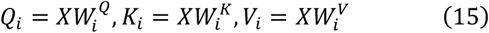

Where 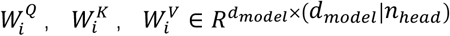 are all learnable matrices. The output of the final head calculation is:

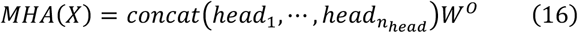

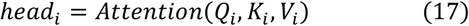

Then, the output of the multi-head attention MHA(X) is added to the original input and normalized, and the resulting output is used as the input to the feed-forward network. For the global encoder, we collect the self-feature, Laplacian vector and WL embedding as input and encoded by the above process, which is shown as follow:

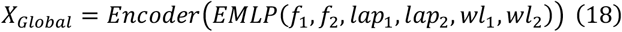

The *Encoder* is a single-layer encoding output. In this study, the number of encoding layers is set to 8. For the local encoder, the neighbor nodes’ representation in subgraph is employed as the network input to focus regional information.

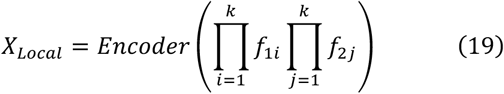

Where operator Π implements the stitching of features into a local matrix. The parament *k* denotes the number of entities in subgraph, feature *f* is the certain row or column vector acquired from similarity matrix *M*_*S*_.

After the encoding process, the encoder output *X* is flattened into a one-dimensional tensor in column. Fully connected layer is then employed to acquire the final prediction output. The binary cross-entropy loss function is adopted for computing the loss, which is defined as follows:

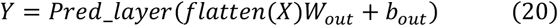

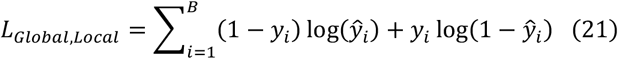

Where *y*_*i*_ is the ground truth association label of the selected nodes, ŷ_*i*_ denotes the predict result which is limited to between 0 and 1, *B* defines the batch size in train strategy. To better balance the training loss between global and local regions, we utilize a hyperparameter *λ* to assign weights to the two above loss functions. Finally, the whole loss function of our model is calculated as follows:

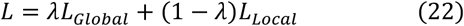

## III. Experiment

In this section, some experiments based on our approach will be implemented to demonstrate its outstanding performance. Firstly, the experiment environment and the main evaluation metrics employed in experiments will be introduced. Then, the result of our model and five other compared models will be shown. Due to the difference in experimental design among these models, we will employ a five-fold cross-validation and use a unified dataset for comparison. In the third part, we visualize the extracted feature vectors to illustrate their effectiveness. After that, various ablation experiments and case studies are designed to confirm the optimality and validity of our method.

### A. Experiment Environment and Evaluation Metrics

In experiment, a single NVIDIA Titan Xp GPU is used, with the neural network framework of pytorch 1.10.1, meanwhile the interpretation is based on Python 3.8. Adam[41] optimizer is employed for overall optimization, with a learning rate set at 1e-3.

A five-fold cross-validation method is employed in the experiment. Specially, the confirmed associated pairs in the current dataset are considered as positive samples, while the molecular pairs with unknown relationships are considered as negative samples. During sampling, we utilize all positive samples and select negative samples in a 1:1 ratio. Afterward, the positive and negative samples are each shuffled and divided into five equal parts. One part is chosen as the testing set alternately and the remaining four parts are regarded as the training set for experimentation. Finally, we average the results to obtain final metric output. The following prediction task metrics are employed as our optimization objectives for the experiment.

(1) AUC: is a wide evaluation metric that measures the performance of a model in terms of its ability to distinguish between positive and negative examples. To obtain the metric, we should calculate the true positive rate (TPR) and false positive rate (FPR) for each probability threshold, which are defined as follows:

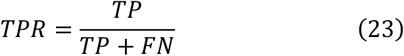

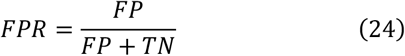

Where, TP and TN are the proportions of instances that are correctly classified as positive or negative by the model. FP and FN are the proportions of instances that are incorrectly classified as positive or negative. Based on the two parameters, we create the ROC (Receiver Operating Characteristic) curve and AUC is the area under the ROC curve.

(2) AUPR: is a commonly used metric to evaluate the performance of classification models, especially in imbalanced dataset. Before the calculation of AUPR, we should know the precision and recall, which are defined as follows:

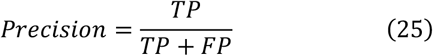

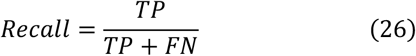

Then, we can calculate the area under the precision-recall curve to obtain AUPR value.

(3) ACC: is a metric to measure the proportion of correct prediction made by the model or classifier out of all the predictions made, which is defined as follows:

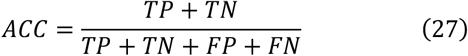

(4) F1: is a measure balancing both precision and recall, which is calculated as the harmonic mean of precision and recall:

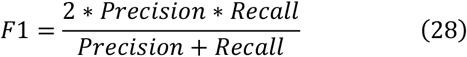

### B. Comparative Evaluation of Performance between Different Methods

To evaluate the effectiveness of our method, we compare it with five other models, including PADLMHOOI[16], CNNMDA[21], DEFLMDA[17], HGCNMDA[42], LDAformer[22]. The PADLMHOOI uses high-order orthogonal iterations to predict potential associations and evaluates predictive performance using leave-one-out cross-validation, which is based on traditional machine learning. CNNMDA and DEFLMDA adopt supervised deep-learning framework for miRNA-disease associations determination. HGCNMDA constructs a heterogeneous network of miRNA, gene, and disease nodes and employs a graph convolution network to predict the associations. LDAformer predicts LncRNA-disease associations by utilizing self-attention mechanism. Then, we compare BertNDA with baselines on the dataset1 and dataset2 under five cross-validation.

As shown in Fig.3, our model achieves state-of-the-art (SOTA) performance in both AUC and AUPR metrics compared to other models, with a 2.0% and 1.7% improvement over the second-best model, respectively. More specific experimental results can be found in the Supplement Table I. After five-fold cross-validation, the evaluation metrics for the tests are as follows: 0.998 for AUC, 0.998 for AUPR, 0.982 for ACC, and 0.982 for F1. These experimental results indicate that our approach based on the Transformer-encoder structure and multi-scale feature fusion has a significant impact on acquiring potential relationships. Moreover, the neighbor representation based on the unconnected subgraph is capable of obtaining multi-hop topological structures effectively, meanwhile the global representation is capable of reflecting the position of the node in the fully graph structure.

**Fig.3.**
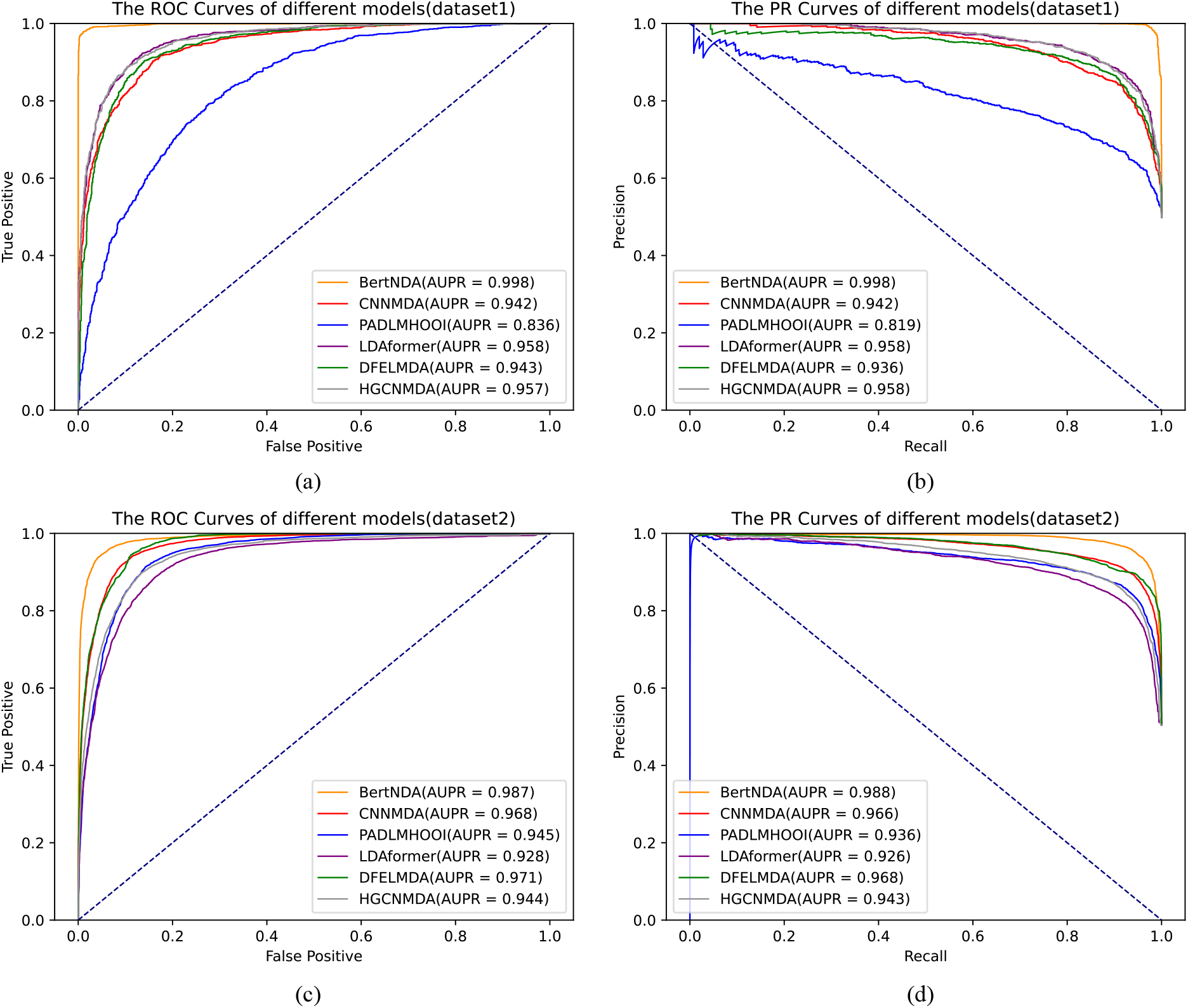
The performance compared with other method. (a)(b) the Curves output in the dataset1; (c)(d) the Curves output in the dataset2.

### C. Visualization of Representation Aggregation and Separation

In this section, we provide visualization to illustrate the impact of features, taking the Laplacian-based feature as an example. As shown in Fig.4, we sample 1000 positive pairs and 1000 negative pairs, and apply t-SNE[43] to reduce the heigher dimensional vectors to two dimensions. The upper plot in each group shows the initial low-dimensional representation, and the bottom plot shows the low-dimensional representation after 20 epochs of training.

**Fig.4.**
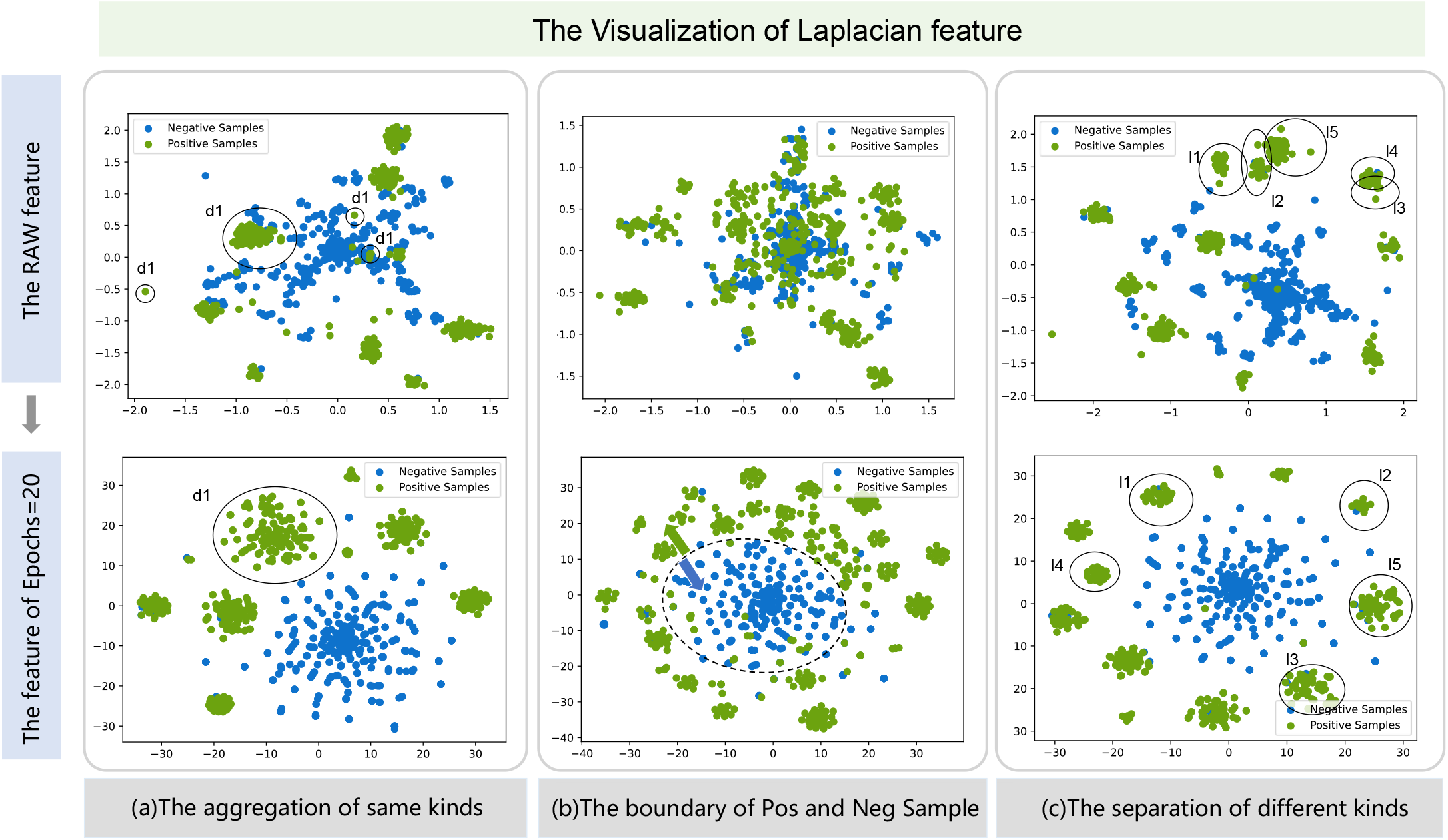
For the results of Laplacian feature extraction, the extraction process uses the t-SNE method, the upper three plots represent the original Laplacian results before training, and the lower three plots represent the Laplacian feature results after training. (a) a characteristic aggregation of a disease; (b) the segmentation boundary of the feature space of positive and negative samples after training; (c) Isolation between different species of lncRNA.

In part (a), the sample pairs are disease-other pairs. It can be observed that the features can aggregate positive pairs that are highly associated with a particular disease (d1: Adenocarcinoma[44]). These pairs with large initial differences are clustered together with the training process, resulting in similar representation. We call this process the “aggregation of same kinds”.

In part (b), the sample pairs are miRNA-other pairs. The result demonstrates that Laplacian features can effectively distinguish positive and negative pairs during training, shown by the clear boundary between the two sets in the bottom plot.

In part (c), the sample pairs are lncRNA-other pairs, and we focus on five lncRNA molecules (l1: CDKN2B-AS1[45], l2: SNHG15[46], l3: ZNRD1-AS1[47], l4: SNHG5[48], l5: DLEU2[49]). Initially, the representations of these pairs are quite similar, which leads to a certain overlap in two-dimensional visualization. However, as training progresses, the potential differences between different lncRNA-other pairs come into being. Concretely, the Euclidean distance between these pairs in the two-dimensional space becomes larger. This indicates that the Laplacian feature can effectively distinguish different molecule pairs and improve the robustness of our prediction.

### D. Ablation studies

#### 1) The Efficient of Different Multi-Scale Representatio

We conduct some ablation experiments to show the impact of specific compositions of the model. To investigate the effectiveness of multi-scale features, we conducted two experiments. First, we explore the influence of the number of nodes in the disconnected subgraph on the results. As shown in Fig.5(a), when we gradually increase the number of adjacent nodes (from 0 to 5), all three metrics (AUC, AUPR, and ACC) improved. Distinctly, ACC increases from 90.3% to over 99.6%(illustrated in Supplement Table 2). This experiment demonstrates that increasing the number of adjacent nodes can effectively improve the local representation of nodes. This is because a greater multi-hop topological structure information can be obtained when there are more nodes, thus allowing the model to better predict potential associations and improving its performance to a certain extent. However, as the number of subgraph nodes continues to increase, the model training time also increases and the model will be overfit. To prevent the occurrence of the above phenomenon, we set the number of adjacent nodes to 6 in the experiment to fully balance prediction performance and time-cost.

**Fig.5.**
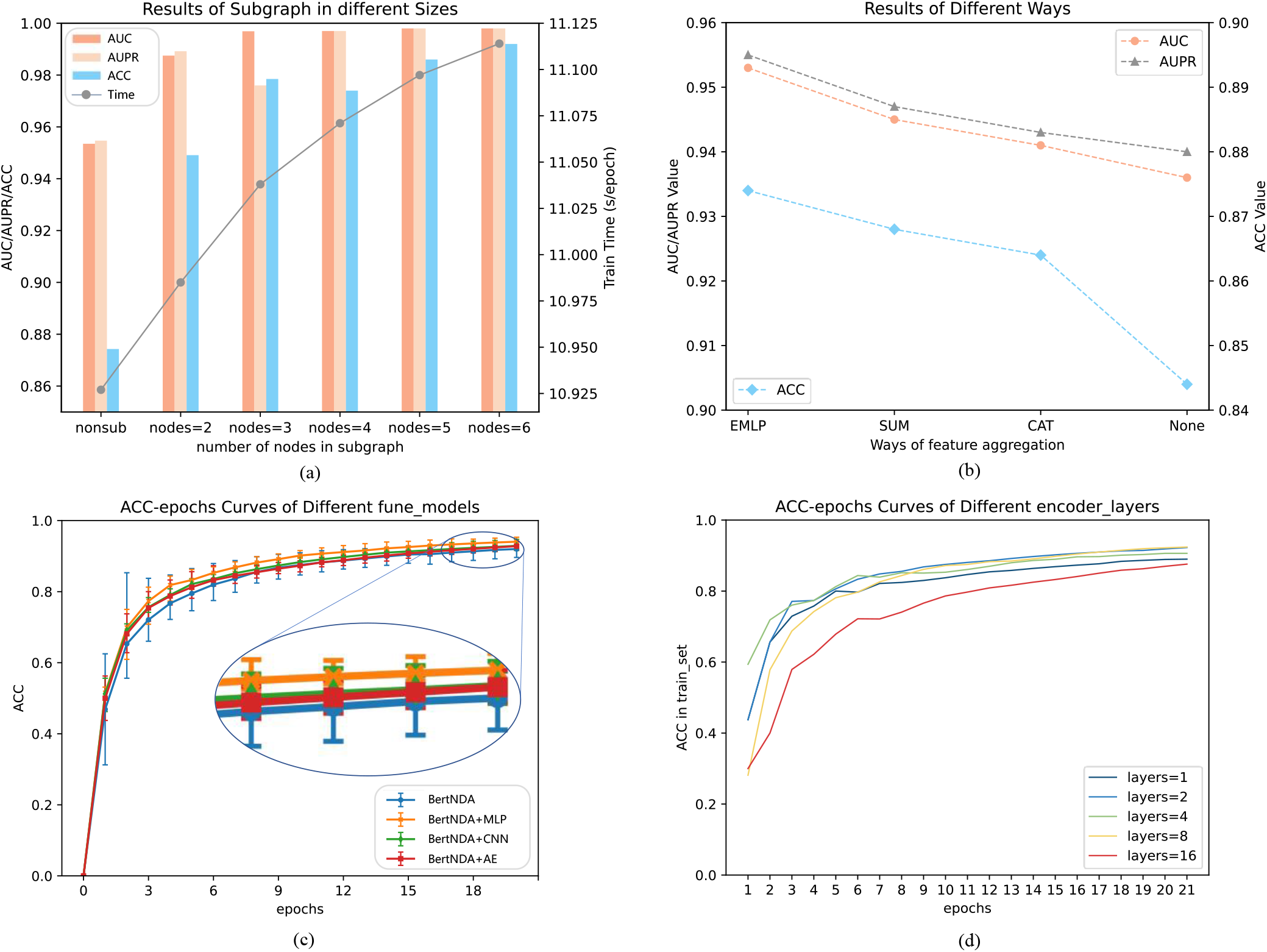
The result of ablation studies. (a) the metrics in multi-size subgraphs; (b) the efficient of EMLP feature combination; (c) the comparison of different fine-tune models; (d) the influence of layers in encoder part.

Then, we explore the impact of different aggregation methods of global representation on our model performance. As shown in Fig.5(b), the aggregation of representation vectors by inserting EMLP can improve the performance of prediction better than the employ of SUM and CAT. Inversely, it will significantly increase the prediction cost of subsequent models and reduce the property of model without any measure.

#### 2) Influence of Model Structure on Prediction Performance

To investigate the influence of model architecture on the prediction performance, we conduct two sets of ablation experiments. Firstly, we add some other fine-tuning structures between the Transformer-encoder backbone network and the original output head to explore their effects, as shown in Fig.5(c). Specifically, we add three types of structures: MLP, CNN[50], and AE[51]. Experimental results show that using MLP leads to a higher performance improvement. We speculate that using CNN or AE would increase the complexity of the overall prediction head. Compared to MLP, achieving the same effect requires longer training time and more training iterations. Therefore, to reduce the training time cost caused by the increased complexity, we prioritize using MLP as the fine-tuning structure for our task.

Finally, the number of encoder layers is often a worthwhile issue to investigate for Transformer serials model. As shown in Fig.5(d), we set different numbers of layers (1, 2, 4, 8, 16) and plot their accuracy curve with respect to epoch. It can be seen that the model converges to better result at a faster rate when the number of layers is 8, meanwhile, this curve has better smoothness than other curves. When we continue to increase the layers to 16, it can be observed that, under the same epoch, the ACC value is lower than other cases. This indicates that with the increase of layers, the global attention can achieve better results, but it also leads to converge at a slower rate. The detailed results of above can be found in the Supplement Table II-V.

#### 3) The Superior Characteristics without Data Leakage

In addition, we also search the impact of result caused by data leakage issues. We find that there is a common problem in the prediction methods of most previous work[16, 21, 42], which is that known association relationships have already been included in the feature extraction process, leading to a certain amount of data leakage in the extracted feature. Meanwhile, considering that we expect to obtain association with unproven but highly probable, we design an ablation experiment to eliminate the data leakage problem. Specifically, we label some known positive sample pairs as negative sample pairs, and then obtained their feature representations based on revised labels. Then, we predict these revised positive sample pairs and other negative sample pairs, and obtained the corresponding evaluation metrics. The results are illustrated in Fig.6. It can be seen that compared to the case without data leakage, all models show a certain degree of decline in the evaluation metrics. Our model’s AUC decreases from 99.7% to 87.3%, and AUPR decreases from 99.4% to 86.9%. However, our model still has SOTA performance. In the five-fold cross-validation, the average and mean of ACC indicators are higher than other models, which is recorded in Supplement Table V for specific.

**Fig.6.**
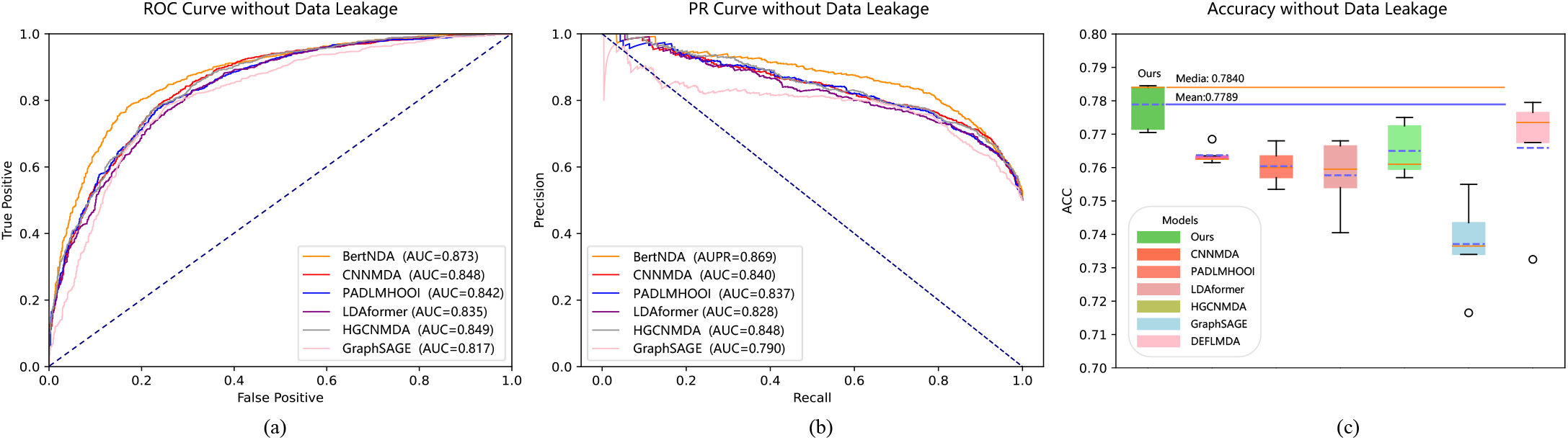
The comparison of experimental results without the data leakage problem. (a) Comparison of ROC Curve and AUC value; (b) Comparison of PR Curve and AUC value; (c) Comparison of ACC acquired by five-fold cross-validation.

### E. Case Studies

To provide a more intuitive demonstration of the training performance of our model, case studies are conducted. Firstly, we use all positive sample pairs, as well as an equal proportion of negative sample pairs sorted by intimacy, as the training set, totaling 14,644 samples. Then, we train the model and combine a certain disease with all other ncRNAs to obtain 520 input pairs. Finally, the predicted values are ranked from small to large and the top 40 molecules are selected for display.

Osteosarcoma is a type of bone cancer that begins in the cells that form bones, which is most often found in the long bones but it can start at any bone[52]. In this case, we select it as our prediction target of disease. The result is shown in Table II, in which most of the associated entities can be found in the original database we chose. While some molecules, such as hsamir-197, NPPAAS1, are not included in our original database but collected in other database, such as MNDR[27], or even couldn’t be found in all current databases. Top 10 of potential miRNAs associated with MALAT1 in dataset1 and top10 of lncRNAs associated with has-mir-214 in dataset2 explored by our model are illustrated in Table III and Table IV, a portion of which can be confirmed in other databases. The result shows that the outputs of our method are effective in discovering potential correlation pairs.

**TABLE I.** The Specifics of Two Datasets

| Dataset | Total associations | miRNA-disease | lncRNA-disease | miRNA-lncRNA | miRNA | lncRNA | disease |
| --- | --- | --- | --- | --- | --- | --- | --- |
| Dataset1 | 7322 | 4802 | 2278 | 242 | 464 | 56 | 121 |
| Dataset2 | 43869 | 23544 | 2038 | 18287 | 1596 | 2189 | 1296 |

**TABLE II.** Top 20 ncRNAS Predicted by Our Model Association with the Osteosarcoma in Dataset1

| Rank | Name | Evidence | Rank | Name | Evidence |
| --- | --- | --- | --- | --- | --- |
| 1 | has-mir-30b | HMDD | 11 | has-mir-204 | HMDD, mircancer |
| 2 | has-mir-335 | HMDD, mircancer | 12 | has-mir-132 | HMDD, mircancer |
| 3 | has-mir-497 | HMDD, mircancer | 13 | has-mir-181c | mircancer |
| 4 | has-mir-192 | HMDD | 14 | has-mir-197 | *MNDR (out of our database)* |
| 5 | has-mir-374a | HMDD, mircancer | 15 | SNHG1 | Lnc2Cancer |
| 6 | has-mir-33b | HMDD, mircancer | 16 | has-mir-377 | HMDD |
| 7 | has-mir-138 | HMDD | 17 | has-let-7a | HMDD, mircancer |
| 8 | has-mir-196a | HMDD | 18 | MIAT | Lnc2Cancer |
| 9 | CCAT1 | Lnc2Cancer | 19 | has-mir-185 | mircancer |
| 10 | hsamir122 | mircancer | 20 | has-mir-449a | HMDD, mircancer |

**TABLE III.** Top 10 miRNAS Predicted by Our Model Associated with the MALAT1 in Dataset1

| Rank | Name | Evidence | Rank | Name | Evidence |
| --- | --- | --- | --- | --- | --- |
| 1 | Has-mir-135b | LncBase v.3 | 6 | Has-mir-421 | LncBase v.3 |
| 2 | Has-mir-19b-3p | LncBase v.3 | 7 | Has-mir-9-5p | LncBase v.3 |
| 3 | Has-mir-4500 | Ago CLIP | 8 | Has-mir-99a | *unconfirmed* |
| 4 | Has-mir-16 | *unconfirmed* | 9 | Has-mir-411-5p | *unconfirmed* |
| 5 | Has-mir-32-5p | LncBase v.3 | 10 | Has-mir-134 | *unconfirmed* |

**TABLE IV.**
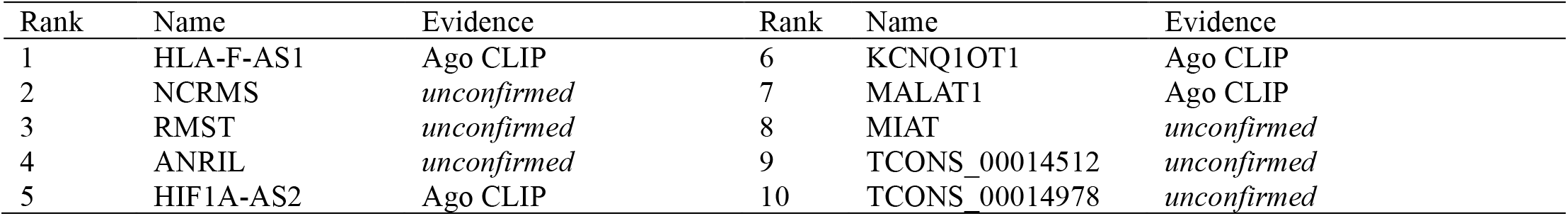
Top 10 lncRNAS Predicted by Our Model Associated with Has-mir-214 in Dataset2

| Rank | Name | Evidence | Rank | Name | Evidence |
| --- | --- | --- | --- | --- | --- |
| 1 | HLA-F-AS1 | Ago CLIP | 6 | KCNQ1OT1 | Ago CLIP |
| 2 | NCRMS | *unconfirmed* | 7 | MALAT1 | Ago CLIP |
| 3 | RMST | *unconfirmed* | 8 | MIAT | *unconfirmed* |
| 4 | ANRIL | *unconfirmed* | 9 | TCONS_00014512 | *unconfirmed* |
| 5 | HIF1A-AS2 | Ago CLIP | 10 | TCONS_00014978 | *unconfirmed* |

### F. Design and Presentation of the Online Platform

We also develop an online platform to showcase our predictive model. This platform enables users to search for molecules of interest and filter the search results. Users can query the relationship of individual molecules through the search box or batch prediction through uploading file. The platform presents the search results in a user-friendly and easy-to-understand manner, offering an intuitive display, and the dataset in our method can also be downloaded. In summary, our platform integrates filtering, querying, predicting, and visualizing to provide users with a versatile and user-friendly website, as depicted in Fig.7.

**Fig.7.**
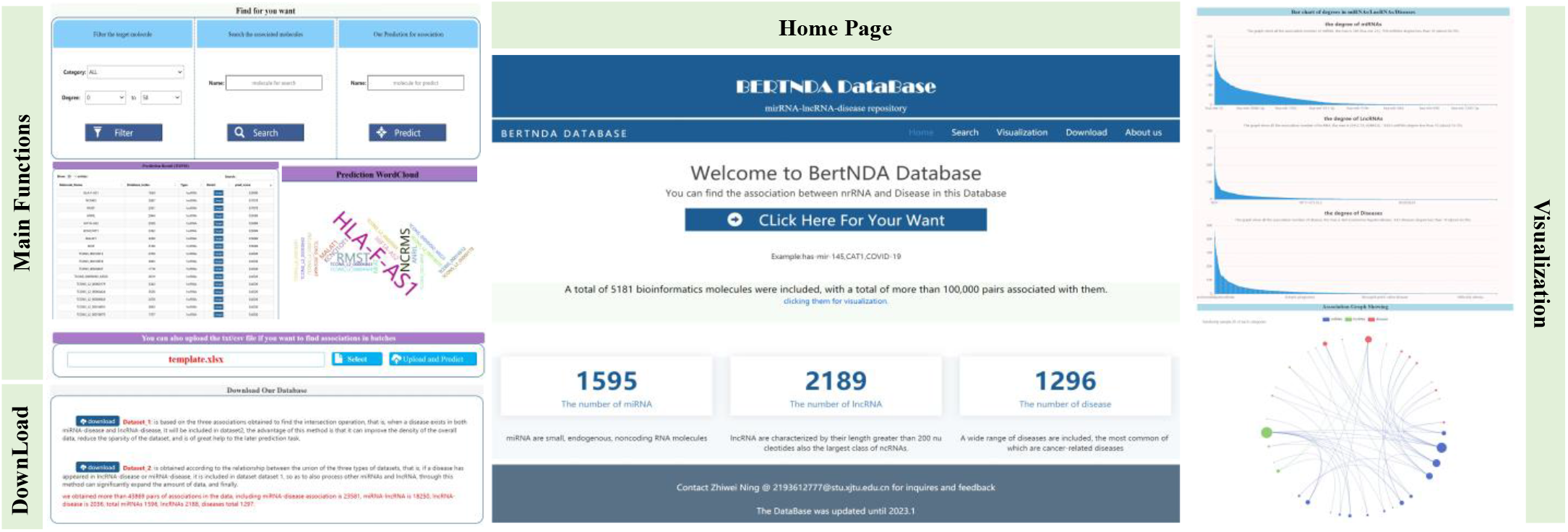
the main pages display in our website, including filter/search/predict functions, download and visualization

## IV. Conclusion and Discussion

Exploring the association between ncRNA and diseases plays a crucial role in disease diagnosis and prevention.

Nevertheless, many existing methods cannot obtain molecular feature representations well from multiple scales and there is little attention on relationships between ncRNAs. Therefore, we propose the BertNDA method, which achieves good prediction results based on the efficient encoding ability of the Transformer encoder model and the powerful representation method of unique information of entities in the graph structure. Through visualization, we introduce the interpretability of the feature expression, and various ablation experiments define the optimal parameters and structure of the model. Finally, we demonstrate the predictive ability of the model for potential molecular pairs through a case study.

However, our model still has some limitations. Firstly, the absolute role encoding method cannot distinguish well between nodes with similar regional structures, which may interfere with the corresponding predicted pairs’ feature representation. Secondly, our method has underutilization of edge information, which may result in the loss of some valid information during feature representation. In the future, edge information can be incorporated into the embedding of the encoder network. Finally, limited by the Transformer structure, as the amount of data increases, the model’s training time will increase rapidly, which is not friendly to the constantly enriched biomedical field. In the future, we can consider appropriately combining with lightweight network structures to achieve better training effects.

### A. Key Points of Our Method

We propose a more comprehensive trilateral prediction relationship between ncRNAs compared with the previous works. Besides, the computational experiments show the superior performance of our method in both data leakage and non-data leakage scenarios.

We employ the heterogeneous graph structure to represent global features extracted from the Laplace matrix and WL absolute position coding, after which a EMLP module is used to fuse them.

To obtain local information of nodes, we sort the neighbor nodes by intimacy and use the unconnected subgraphs to gather information about each node.

An encoder part of the Transformer network is acquired as the backbone network of our method. By extracting the association relationship of nodes in a more macroscopic scale, we improve the effectiveness of the overall prediction.

## Notes

This research was supported by Zhejiang Provincial Natural Science Foundation of China under Grant No. LQ23F020018, Natural Science Basic Research Program of Shaanxi (Program No. 2023-JC-QN-0737), Natural Science Foundation of Sichuan, China (No. 2023NSFSC1416), and National Natural Science Foundation of China (Grant no. 61872288).

Conflict of Interest: none declared.

### Competing Interest Statement

The authors have declared no competing interest.

## References

[1] Bartel D P J C, “MicroRNAs: genomics, biogenesis, mechanism, and function,” Cells, vol. 116, no. 2, pp. 281–297, 2004.

[2] Iyer M K, et al., “The landscape of long noncoding RNAs in the human transcriptome,” Nature Genetics, vol. 47, no. 3, pp. 199–208, 2015.

[3] Managadze D, et al., “Negative correlation between expression level and evolutionary rate of long intergenic noncoding RNAs,” Genome biology and evolution, vol. 3, pp. 1390–1404, 2011.

[4] Mattick J S, “The genetic signatures of noncoding RNAs,” PLoS genetics, vol. 5, no. 4, pp. e1000459, 2009.

[5] Yang Z, et al., “Overexpression of long non-coding RNA HOTAIR predicts tumor recurrence in hepatocellular carcinoma patients following liver transplantation,” Annals of surgical oncology, vol. 18, pp. 1243–1250, 2011.

[6] Zhang Z, et al., “Evaluation of novel gene UCA1 as a tumor biomarker for the detection of bladder cancer,” Zhonghua yi xue za zhi, vol. 92, no. 6, pp. 384–387, 2012.

[7] Ambros V J N, “The functions of animal microRNAs,” Nature, vol. 431, no. 7006, pp. 350–355, 2004.

[8] Karp X, Ambros V, “Encountering microRNAs in cell fate signaling,” Science, vol. 310, no. 5752, pp. 1288–1289, 2005.

[9] Cheng A M, et al., “Antisense inhibition of human miRNAs and indications for an involvement of miRNA in cell growth and apoptosis,” Nucleic Acids Research, vol. 33, no. 4, pp. 1290–1297, 2005.

[10] Miska E A, “How microRNAs control cell division, differentiation and death,” Current opinion in genetics & development, vol. 15, no. 5, pp. 563–568, 2005.

[11] Paraskevopoulou M D, Hatzigeorgiou A G J L N-C R M, Protocols, “Analyzing miRNA–lncRNA interactions,” Methods in Molecular Biology, pp. 271–286, 2016.

[12] Li N, et al., “The role of microRNA and LncRNA–MicroRNA interactions in regulating ischemic heart disease,” Cardiovascular Pharmacology and Therapeutics, vol. 22, no. 2, pp. 105–111, 2017.

[13] Ling H, et al., “Junk DNA and the long non-coding RNA twist in cancer genetics,” Oncogene, vol. 34, no. 39, pp. 5003–5011, 2015.

[14] Bandyopadhyay S, et al., “Development of the human cancer microRNA network,” Silence, vol. 1, no. 1, pp. 1–14, 2010.

[15] Lu M, et al., “An analysis of human microRNA and disease associations,” PloS one, vol. 3, no. 10, pp. e3420, 2008.

[16] Xuan Z, et al., “A novel method for predicting disease-associated LncRNA-MiRNA pairs based on the higher-order orthogonal iteration,” Computational and mathematical methods in medicine, 2019.

[17] Yu L, Zheng Y, Gao L J B I B, “MiRNA–disease association prediction based on meta-paths,” Briefings in Bioinformatics, vol. 23, no. 2, pp. 2022.

[18] Chen X, Yan G-Y J B, “Novel human lncRNA–disease association inference based on lncRNA expression profiles,” Bioinformatics, vol. 29, no. 20, pp. 2617–2624, 2013.

[19] Xuan P, et al., “Graph convolutional network and convolutional neural network based method for predicting lncRNA-disease associations,” Cells, vol. 8, no. 9, pp. 1012, 2019.

[20] Li G, et al., “Using Graph Attention Network and Graph Convolutional Network to Explore Human CircRNA–Disease Associations Based on Multi-Source Data,” Frontiers in Genetics, vol. 13, pp. 64, 2022.

[21] Peng J, et al., “A learning-based framework for miRNA-disease association identification using neural networks,” Bioinformatics, vol. 35, no. 21, pp. 4364–4371, 2019.

[22] Zhou Y, et al., “LDAformer: predicting lncRNA-disease associations based on topological feature extraction and Transformer encoder,” Briefings in Bioinformatics, vol. 23, no. 6, pp. 2022.

[23] Zhang J, et al., “Graph-bert: Only attention is needed for learning graph representations,” arXiv preprint 2001.05140, vol. no. pp. 2020.

[24] Kozomara A, Griffiths-Jones S J N a R, “miRBase: integrating microRNA annotation and deep-sequencing data,” Nucleic Acids Research, vol. 39, pp. D152–D157, 2010.

[25] Li Y, et al., “HMDD v2. 0: a database for experimentally supported human microRNA and disease associations,” Nucleic Acids Research, vol. 42, no. D1, pp. D1070–D1074, 2014.

[26] Jiang Q, et al., “miR2Disease: a manually curated database for microRNA deregulation in human disease,” Nucleic Acids Research, vol. 37, pp. D98–D104, 2009.

[27] Ning L, et al., “MNDR v3. 0: mammal ncRNA–disease repository with increased coverage and annotation,” Nucleic Acids Research, vol. 49, no. D1, pp. D160-D164, 2021.

[28] Bao Z, et al., “LncRNADisease 2.0: an updated database of long noncoding RNA-associated diseases,” Nucleic Acids Research, vol. 47, no. D1, pp. D1034–D1037, 2019.

[29] Xie B, et al., “miRCancer: a microRNA–cancer association database constructed by text mining on literature,” Bioinformatics, vol. 29, no. 5, pp. 638–644, 2013.

[30] Gao Y, et al., “Lnc2Cancer 3.0: an updated resource for experimentally supported lncRNA/circRNA cancer associations and web tools based on RNA-seq and scRNA-seq data,” Nucleic Acids Research, vol. 49, no. D1, pp. D1251–D1258, 2021.

[31] Li J-H, et al., “starBase v2. 0: decoding miRNA-ceRNA, miRNA-ncRNA and protein–RNA interaction networks from large-scale CLIP-Seq data,” Nucleic Acids Research, vol. 42, no. D1, pp. D92–D97, 2014.

[32] Lowe H J, Barnett G O J J, “Understanding and using the medical subject headings (MeSH) vocabulary to perform literature searches,” Jama, vol. 271, no. 14, pp. 1103–1108, 1994.

[33] Wang D, et al., “Inferring the human microRNA functional similarity and functional network based on microRNA-associated diseases,” Bioinformatics, vol. 26, no. 13, pp. 1644–1650, 2010.

[34] Murphy R, et al., “Relational pooling for graph representations,” in International Conference on Machine Learning. 2019, pp. 4663–4673

[35] Velickovic P, et al., “Graph attention networks,” stat, vol. 1050, no. 20, pp. 10.48550, 2017.

[36] Dwivedi V P, et al., “Benchmarking graph neural networks,” Journal of machine learning research, 2020.

[37] Niepert M, Ahmed M, Kutzkov K, “Learning convolutional neural networks for graphs,” in International conference on machine learning. 2016, pp. 2014–2023

[38] Vaswani A, et al., “Attention is all you need,” Advances in neural information processing systems, vol. 30, pp. 2017.

[39] Xie Q, et al., “Venet: Voting enhancement network for 3d object detection,” in Proceedings of the IEEE/CVF International Conference on Computer Vision. 2021, pp. 3712–3721

[40] Adamic E, Adar L A, “Friends and neighbors on the web,” Social Network Analysis, vol. 25, no. 3, pp. 211–230, 2003.

[41] Kingma D P, Ba J J a P A, “Adam: A method for stochastic optimization,” arXiv preprint 1412.6980, 2014.

[42] Peng W, et al., “Predicting miRNA-disease associations from miRNA-gene-disease heterogeneous network with multi-relational graph convolutional network model,” IEEE/ACM Transactions on Computational Biology and Bioinformatics, 2022.

[43] Van Der Maaten L, Hinton G J J O M L R, “Visualizing data using t-SNE,” Journal of machine learning research, vol. 9, no. 11, 2008.

[44] Khosla D, et al., “Small bowel adenocarcinoma: An overview,” World Journal of Gastrointestinal Oncology, vol. 14, no. 2, pp. 413, 2022.

[45] Song C, et al., “CDKN2B-AS1: an indispensable long non-coding RNA in multiple diseases,” Current Pharmaceutical Design, vol. 26, no. 41, pp. 5335–5346, 2020.

[46] Saeinasab M, et al., “SNHG15 is a bifunctional MYC-regulated noncoding locus encoding a lncRNA that promotes cell proliferation, invasion and drug resistance in colorectal cancer by interacting with AIF,” Experimental & Clinical Cancer Research, vol. 38, pp. 1–16, 2019.

[47] Wang J, et al., “lncRNA ZNRD1-AS1 promotes malignant lung cell proliferation, migration, and angiogenesis via the miR-942/TNS1 axis and is positively regulated by the m6A reader YTHDC2,” Molecular Cancer, vol. 21, no. 1, pp. 1–19, 2022.

[48] Li Y-H, et al., “LncRNA SNHG5: a new budding star in human cancers,” Gene, vol. 749, pp. 144724, 2020.

[49] Xu W, et al., “DLEU2: a meaningful long noncoding RNA in oncogenesis,” Current Pharmaceutical Design, vol. 27, no. 20, pp. 2337–2343, 2021.

[50] Krizhevsky A, Sutskever I, Hinton G E J C O T A, “Imagenet classification with deep convolutional neural networks,” Communications of the ACM, vol. 60, no. 6, pp. 84–90, 2017.

[51] Michelucci U J a P A, “An introduction to autoencoders,” arXiv preprint 2201.03898, 2022.

[52] Beird H C, et al., “Osteosarcoma,” Nature Reviews Disease Primers, vol. 8, no. 1, pp. 77, 2022.

