## Supplement for "BertNDA: A Model Based on Graph-Bert and Multi-scale Information Fusion for ncRNA-disease Association Prediction"

### Supplementary Materials

TABLE I

THE RESULTS COMPARED WITH OTHER BASELINE MODELS IN DATASET1

| Metrics | Methods | | | | | |
|---|---|---|---|---|---|---|
| | Our | CNNMDA | PADLMHOOI | LDAformer | DFELMDA | HGCNMDA |
| AUC | **99.83** | 94.17 | 83.62 | 95.82 | 94.33 | 95.71 |
| AUPR | **99.84** | 94.21 | 81.92 | 95.83 | 93.60 | 95.78 |
| ACC | **98.19** | 86.04 | 75.42 | 89.11 | 87.77 | 88.80 |
| F1 | **98.18** | 85.64 | 75.09 | 89.05 | 87.59 | 88.56 |

THE RESULTS COMPARED WITH OTHER BASELINE MODELS IN DATASET2

| Metrics | Methods | | | | | |
|---|---|---|---|---|---|---|
| | Our | CNNMDA | PADLMHOOI | LDAformer | DFELMDA | HGCNMDA |
| AUC | **98.70** | 96.82 | 94.47 | 92.84 | 97.10 | 94.38 |
| AUPR | **98.78** | 96.56 | 93.61 | 92.61 | 96.75 | 94.27 |
| ACC | **93.43** | 90.87 | 87.63 | 85.90 | 90.09 | 87.85 |
| F1 | **93.21** | 90.83 | 87.58 | 86.06 | 89.97 | 87.92 |

TABLE II

THE INFLUENCE OF THE NODE SIZES IN SUBGRAPH

| Metrics | Number of nodes in subgraph | | | | | |
|---|---|---|---|---|---|---|
| | nodes=1 | nodes=2 | nodes=3 | nodes=4 | nodes=5 | nodes=6 |
| Time cost (s/epoch) | 10.927 | 10.985 | 11.038 | 11.071 | 11.097 | 11.114 |
| AUC | 95.36 | 98.75 | 99.68 | 99.75 | 99.91 | **99.97** |
| AUPR | 95.47 | 98.92 | 99.72 | 99.79 | 99.91 | **99.97** |
| ACC | 87.44 | 94.91 | 97.84 | 97.43 | 98.63 | **99.28** |
| F1 | 86.93 | 94.86 | 97.83 | 97.46 | 98.62 | **99.28** |

TABLE III

THE RESULTS OF DIFFERENT FEATURE AGGREGATION METHODS

| Metrics | Feature aggregation way | | | |
|---|---|---|---|---|
| | E(element-weight) MLP | Weight-multiply and sum up | Concatenate and using MLP | None process of feature aggregation |
| AUC | **95.36** | 94.59 | 94.15 | 93.64 |
| AUPR | **95.47** | 94.73 | 94.33 | 94.02 |
| ACC | **87.44** | 86.89 | 86.41 | 84.50 |
| F1 | 86.93 | **87.29** | 86.72 | 82.74 |

TABLE IV

THE ACC OF DIFFERENT FINE-TUNE MODEL

| Metric | Different fine-tune models | | | |
|---|---|---|---|---|
| | Origin BertNDA | BertNDA+MLP | BertNDA+CNN | BertNDA+AE |
| ACC | $91.99^{+1.99}_{-2.38}$ | $\mathbf{94.06^{+1.25}_{-0.78}}$ | $92.91^{+1.81}_{-1.31}$ | $92.81^{+1.25}_{-1.09}$ |

TABLE V

THE AVERAGE ACC OF DIFFERENT SIZE OF LAYERS IN ENCODER NETWORK IN EACH EPOCH (1-20):

| ACC | The number of layers | | | | |
|---|---|---|---|---|---|
| | Layers=1 | Layers=2 | Layers=4 | Layers=8 | Layers=16 |
| epoch=1 | 0.4375 | 0.4375 | **0.5938** | 0.2813 | 0.3000 |
| epoch=2 | 0.6563 | 0.6563 | **0.7188** | 0.5781 | 0.4000 |
| epoch=3 | 0.7292 | **0.7708** | 0.7604 | 0.6875 | 0.5792 |
| epoch=4 | 0.7578 | **0.7734** | **0.7734** | 0.7422 | 0.6219 |
| epoch=5 | 0.8000 | 0.8063 | **0.8125** | 0.7813 | 0.6788 |
| epoch=6 | 0.7969 | 0.8333 | **0.8438** | 0.7969 | 0.7219 |
| epoch=7 | 0.8214 | **0.8482** | 0.8393 | 0.8259 | 0.7214 |
| epoch=8 | 0.8242 | **0.8555** | 0.8516 | 0.8438 | 0.7406 |
| epoch=9 | 0.8299 | **0.8681** | 0.8507 | 0.8611 | 0.7660 |
| epoch=10 | 0.8375 | **0.8750** | 0.8531 | 0.8719 | 0.7863 |
| epoch=11 | 0.8466 | **0.8807** | 0.8608 | 0.8750 | 0.7972 |
| epoch=12 | 0.8542 | **0.8854** | 0.8698 | 0.8828 | 0.8089 |
| epoch=13 | 0.8582 | **0.8918** | 0.8798 | 0.8846 | 0.8164 |
| epoch=14 | 0.8639 | **0.8973** | 0.8862 | 0.8906 | 0.8250 |
| epoch=15 | 0.8688 | **0.9021** | 0.8896 | 0.8979 | 0.8325 |
| epoch=16 | 0.8731 | **0.9063** | 0.8965 | 0.9043 | 0.8410 |
| epoch=17 | 0.8768 | **0.9099** | 0.8971 | **0.9099** | 0.8504 |
| epoch=18 | 0.8837 | 0.9132 | 0.9010 | **0.9149** | 0.8587 |
| epoch=19 | 0.8865 | 0.9145 | 0.9030 | **0.9194** | 0.8628 |
| epoch=20 | 0.8891 | 0.9188 | 0.9063 | **0.9219** | 0.8697 |

TABLE VI

THE AUC/AUPR OF DIFFERENT MODELS WITHOUT DATA LEAKAGE:

| Metrics | models | | | | | |
|---|---|---|---|---|---|---|
| | Ours | CNNMDA | PADLMHOOI | LDAformer | HGCNMDA | GraphSAGE |
| AUC | **87.3** | 84.8 | 84.2 | 83.5 | 84.9 | 81.7 |
| AUPR | **86.9** | 84.0 | 83.7 | 82.8 | 84.8 | 79.0 |

THE ACC OF FIVE-FOLD CROSS-VALIDATION WITHOUT DATA LEAKAGE

| Models | ACC value of different folds | | | | | |
|---|---|---|---|---|---|---|
| | Fold_0 | Fold_1 | Fold_2 | Fold_3 | Fold_4 | Average |
| Ours | **78.45** | **78.40** | **77.05** | 77.15 | **78.40** | **77.89** |
| CNNMDA | 76.15 | 76.35 | 76.85 | 76.25 | 76.25 | 76.37 |
| PADLMHOOI | 76.00 | 76.80 | 75.35 | 75.70 | 76.35 | 76.04 |
| LDAformer | 76.80 | 74.05 | 75.95 | 75.40 | 76.65 | 75.77 |
| HGCNMDA | 77.50 | 77.25 | 75.70 | 75.95 | 76.10 | 76.50 |
| GraphSAGE | 73.40 | 75.50 | 74.35 | 73.65 | 71.65 | 73.71 |
| DEFLMDA | 77.35 | 76.75 | 73.25 | **77.95** | 77.65 | 76.59 |

#### HEATMAP

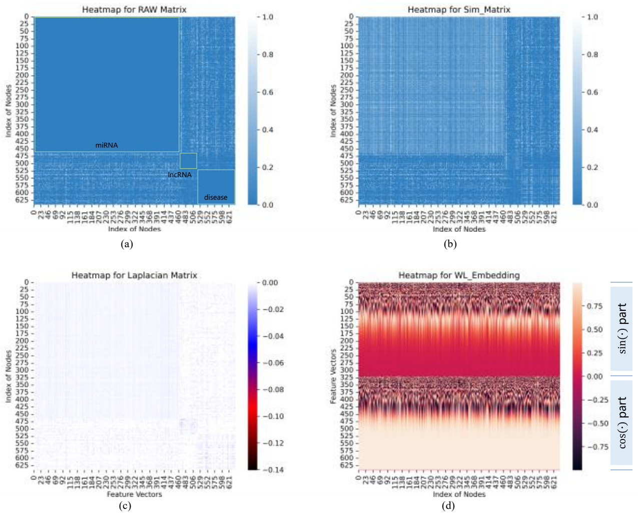


Fig.1. The heatmap of different representation matrix: (a) the raw association matrix; (b) the similarity matrix; (c) Laplacian matrix; (d) WL-embedding matrix in dataset1, the dimension equals to 641.

#### THE INTRODUCTION OF ONLINE-PLATFORM WEBSITE

We design an online display platform for this work, which enables users to filter for certain molecule or search for molecules of interest. Users can query the relationship of individual molecules through the search box or predict in batch through uploading file. The platform presents the search results in an easy-to-understand manner, offering an intuitive display, and the datasets in our method can also be downloaded. In summary, our platform integrates filtering, querying, predicting, and visualizing to provide users with a versatile and user-friendly website.

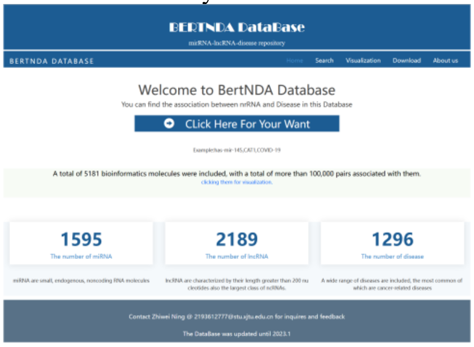


Fig.2. Home Page of online platform, which shows the summary introduction of our dataset and mean function

In Fig.2 the “Home” page brief information about this website is presented, and the page can jump to the "Search" or "Visualization" position by clicking the corresponding hyperlink.

In “Search” page, users can filter miRNA, lncRNA and disease category in the “Filter the target molecule” column, or further subdivide the scale by selecting the degree of the appropriate interval based on the strength of the existing correlation, which is illustrated in Fig.3.

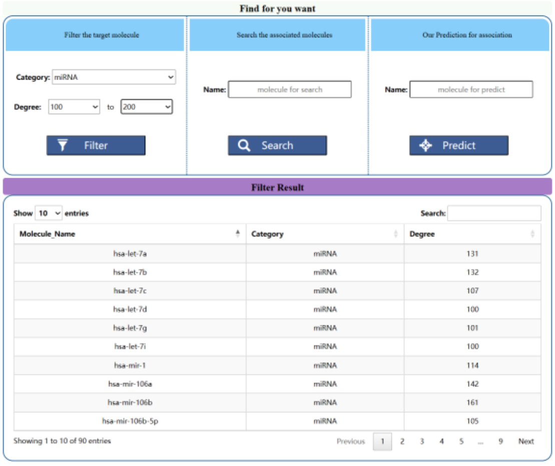


Fig.3. Filter result of category in “miRNA” and degree from “100” to “200”, and total 90 entities are found.

After choosing the target entity, all the association relationships about the molecules or diseases in this dataset can be displayed through clicking “Search” button. Then click “Detail” for specific information, where the disease information is mainly from “Malacard”, the data source of miRNA is from “miRBase” and the lncRNA is from “ENSEMBL” and “NCBI” (due to the inadequacy of the these data sources, it may lead to the loss of specific information for some molecules, which mainly occurs in lncRNA). The concrete content is shown in Fig.4.

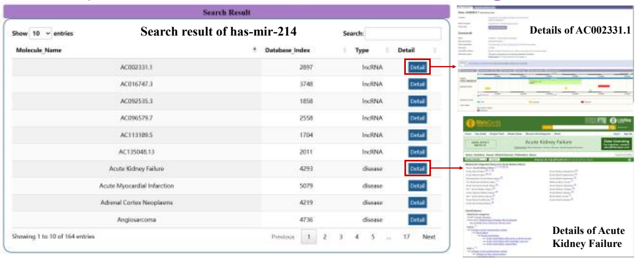


Fig.4. Search result of has-mir-214 associated lncRNAs or diseases in our dataset2. Right part illustrates the details of certain entity.

Furthermore, swe also employ the model to make potential association predictions on input entities. When clicking "Predict", the related molecules will be predicted and displayed. “Pred_score” represents the normalized prediction score and we only show the top 20 results here(illustrated in Fig.5).

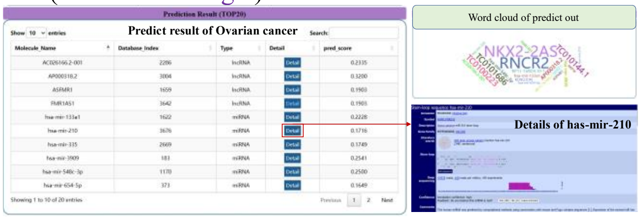


Fig.5. Prediction results of “Ovarian cancer” through our model. Right part displays the word cloud and details.

If users need to predict various molecules at same time, the task can be submitted by uploading the input file (supporting docx, xlsx, txt file formats). After uploading process, the prediction of TOP20 potential molecules related with each input entity will be ac quired and output .xlsx file will be downloaded automatically (shown in Fig.6).

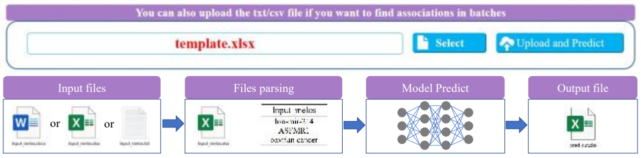


Fig.6. Flow chart of Potential association prediction by uploading file, including upload, parse, predict and output composition.

Taking an example, the information of the output file is employed as follows (the entities included in upload file are: has-mir-214, ASFMR1 and Ovarian cancer).

TABLE VII
THE PART PREDICTION RESULT OF FILE SUBMISSION

| Input_entities | Pred_entities | Pred_categories | Pred_scores |
|---|---|---|---|
| hsa-mir-214 | HLA-F-AS1 | lncRNA | 0.999 |
| | TCONS_L2_00018070 | lncRNA | 0.4526 |
| | TCONS_00024647 | lncRNA | 0.4526 |
| | HIF1A-AS2 | lncRNA | 0.5089 |
| | RMST | lncRNA | 0.7078 |
| | TCONS_L2_00020565 | lncRNA | 0.4526 |
| | MIAT | lncRNA | 0.5089 |
| | KCNQ1OT1 | lncRNA | 0.5089 |
| | TCONS_L2_00006843 | lncRNA | 0.4526 |
| | TCONS_L2_00021262 | lncRNA | 0.4526 |
| | TCONS_00014512 | lncRNA | 0.4526 |
| | TCONS_00090092_MEG3 | lncRNA | 0.4526 |
| | ANRIL | lncRNA | 0.5089 |
| | TCONS_00014978 | lncRNA | 0.4526 |
| | TCONS_L2_00000179 | lncRNA | 0.4526 |
| ASFMR | hsa-mir-134-5p | miRNA | 0.0348 |
| | hsa-mir-134 | miRNA | 0.0216 |
| | hsa-mir-424 | miRNA | 0.0074 |
| | hsa-mir-510 | miRNA | 0.1688 |
| | hsa-mir-130b | miRNA | 0.0157 |
| | hsa-mir-1237 | miRNA | 0.007 |
| | hsa-mir-574 | miRNA | 0.0529 |
| | Neurodevelopmental Disorder | disease | 0.999 |
| | Carcinoma Ovarian Serous | disease | 0.0093 |
| | Spinal Chordoma | disease | 0.0167 |
| | Central Nervous System | disease | 0.0294 |
| | Skin Hemangioma | disease | 0.0061 |
| | Neurological | disease | 0.2321 |
| ovarian cancer | hsa-mir-3909 | miRNA | 0.2541 |
| | hsa-mir-654-5p | miRNA | 0.1649 |
| | hsa-mir-548c-3p | miRNA | 0.25 |
| | hsa-mir-133a1 | miRNA | 0.2228 |
| | ASFMR1 | lncRNA | 0.1903 |
| | RP11-169D4.10-1 | lncRNA | 0.2335 |
| | TC0101686 | lncRNA | 0.5124 |
| | AC026166.2-001 | lncRNA | 0.2335 |
| | RRP1B | lncRNA | 0.1726 |
| | MIR21 | lncRNA | 0.2335 |
| | RP11-161M6.2 | lncRNA | 0.1879 |
| | KCNQ1DN | lncRNA | 0.1993 |
| | hsa-mir-335 | miRNA | 0.1749 |
| | RNCR2 | lncRNA | 0.999 |
| | NKX2-2AS | lncRNA | 0.999 |
